# FASTKD5 processes mitochondrial pre-mRNAs at non-canonical cleavage sites

**DOI:** 10.1101/2024.07.18.603998

**Authors:** Hana Antonicka, Woranontee Weraarpachai, Ana Vučković, Hauke S. Hillen, Eric A. Shoubridge

## Abstract

The regulation of mammalian mitochondrial gene expression is largely post-transcriptional and the first step in translating the 13 polypeptides encoded in mtDNA is endonucleolytic cleavage of the primary polycistronic transcripts. As the rRNAs and most of the mRNAs in mtDNA are flanked by tRNAs, the release of the mature RNAs occurs mostly by excision of the tRNAs. Processing the non-canonical mRNAs, not flanked by tRNAs, requires FASTKD5, but the molecular mechanism remains unknown. To investigate this, we created and characterized a knockout cell line to use as an assay system. The absence of FASTKD5 resulted in a severe combined OXPHOS assembly defect due to the inability to translate mRNAs with unprocessed 5’-UTRs. Analysis of RNA processing of FASTKD5 variants allowed us to map amino acid residues essential for function. Remarkably, this map was RNA substrate-specific, arguing against a one size fits all model. A reconstituted *in vitro* system with purified FASTKD5 protein and synthetic RNA substrates showed that FASTKD5 on its own was able to cleave client substrates correctly, but not non-specific RNA sequences. These results establish FASTKD5 as the missing piece of the biochemical machinery required to completely process the primary mitochondrial transcript.

## Introduction

Mammalian mitochondrial DNA (mtDNA) codes for 13 polypeptides that are essential structural subunits of the oxidative phosphorylation (OXPHOS) complexes, and the 22 tRNAs and 2 rRNAs required for their translation on specialized ribosomes in the mitochondrial matrix. Transcription of mtDNA produces two polycistronic transcripts from the so-called heavy and light strands of mtDNA that must be processed by endoribonucleases to release the mature RNA species. Maturation of most mitochondrial RNAs occurs by cleavage of tRNAs at the 5’ end by mitochondrial RNase P and at the 3’ end by mitochondrial RNase Z, a concept known for decades as the tRNA punctuation model^1^. Mitochondrial RNase P lacks an RNA component and is rather composed of three protein subunits ^2^. A cryo-EM structure of the enzyme bound to RNA showed that two of the subunits (TRMT10C, SDR5C1) form a platform that binds RNA, positioning the catalytic subunit PRORP to cleave the tRNA substrate at the precise 5’end ^3^. ELAC2, the RNase Z that processes tRNA 3’ends, is targeted to both the nucleus and mitochondria ^4^ and recent cryo-EM structures show that the same accessory subunits used by RNase P are also required for the complete 3’end processing of mitochondrial tRNAs with degenerate structures ^5–7^.

The molecular machinery required to mature mRNAs that are not flanked by tRNAs remained unknown until we demonstrated that depletion of FASTKD5, a member of the FASTK-family of proteins, resulted in the accumulation of unprocessed transcripts from precisely those non-canonical open reading frames: the 5’ end of cytochrome *c* oxidase subunit 1 (*CO1*), the junction between the bicistronic *ATP8/6* mRNA and *CO3*, and the junction between the *ND5* and *cytb* mRNAs ^8^. These results were subsequently confirmed in another cell line^9^. It is not known whether FASTKD5 complexes with or recruits other proteins, or whether it possesses endoribonuclease activity itself.

The FASTK (Fas-activated serine-threonine kinase) family is comprised of 6 proteins: FASTK and FASTKD1-5. The FASTK family appeared early in the evolution of Metazoa and phylogenetic analysis shows that expansion of the family occurred in the vertebrate lineage. FASTK has a cytosolic and mitochondrial isoform, but all others localize exclusively to the mitochondrial matrix (reviewed in ^10^). While all FASTK family members appear to have specific roles in mitochondrial RNA transactions, the precise molecular mechanisms of action remain largely obscure^9^. FASTK, FASTKD1, FASTKD2, and FASTKD5 co-localize with markers of mitochondrial RNA granules, which are centers of posttranscriptional processing and modification of mitochondrial RNAs ^8^, the others appear more broadly distributed in the matrix. All contain a predicted mitochondrial matrix targeting sequence (MTS), two FAST domains (FAST1, FAST2, of unknown function), and a so-called RAP domain (RNA-binding domain enriched in apicomplexans) at the extreme C-terminus. The RAP domain of FASTKD4 was suggested to adopt an endonuclease fold similar to that found in PD-(D/E)-XK nucleases ^11^, but such an activity has never been demonstrated. All mammalian FASTK family members also contain a variable number of heptatricopeptide (HPR) repeats which overlap with the FAST1 domain. These repeats are reminiscent of octotricopeptide repeat domains (OPR) ^12^ and the PPR domains of pentatricopeptide repeat proteins ^13^, both of which are RNA-binding domains.

Here we have investigated the biochemical consequences of deletion of *FASTKD5*, mapped the amino acid residues essential for function using a null cell line as an assay system, and demonstrated, using an *in vitro* reconstituted system, that FASTKD5 on its own has can process specific pre-mRNA substrates.

## Materials and Methods

### Key reagents

Supplementary table (**Table S1**) presents a list of antibodies, a recombinant protein and recombinant DNAs, cell lines, oligonucleotides, and oligoribonucleotides used in this study.

### Cell lines

The 143B cell line, derived knock-out (KO) clones and derived cell lines expressing FASTKD5 constructs were grown in high glucose Dulbecco’s modified Eagle medium (DMEM) supplemented with 10% fetal bovine serum, at 37°C in an atmosphere of 5% CO_2_. The medium for the knock-out clones and derived cell lines expressing FASTKD5 constructs was supplemented with 50 µg/ml of uridine. Cell lines were routinely tested for mycoplasma contamination.

For protein stability experiments, selected cell lines were grown in 6-well plates in regular medium supplemented with 10 µg/ml cycloheximide for 0, 4, 8, 16, 24 and 32 hours, after which cells were collected by trypsinization for downstream analysis.

### Generation of knock-out and overexpression cell lines

Knock-out cell lines of FASTKD5 were generated by CRISPR/Cas9–mediated gene editing in 143B cells using a gene-specific target sequence (sgKD5 sequence is indicated in **Table S1**) cloned into pSpCas9(BB)-2A-Puro (PX459) V2.0 ^14^. The plasmid was transfected into 143B cells using Lipofectamine 3000 (Thermo Fisher Scientific) according to the manufacturer’s instructions. The following day, transfected cells were selected by addition of puromycin (2.5 μg/ml) for 2 days. Single cell clones were screened for loss of FASTKD5 protein by immunoblotting with a FASTKD5 antibody (Sigma, **Table S1**) and frameshift mutations were confirmed by genomic sequencing. Two clones were selected for downstream analysis.

143B and the KO cell lines were reconstituted with FASTKD5 constructs by retroviral infection with a virus produced in Phoenix cells transfected with pBABE-3xFLAG-Puro plasmids as described previously ^15^. Briefly, retroviral constructs were transiently transfected into the Phoenix packaging cell line using the HBS/Ca_3_(PO_4_)_2_ method. 143B and KO cells were infected 48 hours later by exposure to virus-containing medium in the presence of 4 μg/ml of polybrene as described ^16^.

### Plasmids

The wild-type FASTKD5 cDNA in pDONR221 ^17^ was cloned into Gateway modified pBABE-3xFLAG-Puro plasmid ^18^ using Gateway technology (Thermo Fisher Scientific). The single point mutations and FASTKD5 deletion mutants were engineered using QuikChange Lightning Site-Directed Mutagenesis Kit (Agilent) with primers indicated in **Table S1** according to manufacturer’s instructions.

For recombinant protein preparation, the sequence encoding FASTKD5 lacking the N-terminal mitochondrial targeting sequence (Δ1–27) was PCR amplified from human cDNA obtained from B-lymphocytes. It was cloned into the 438-C vector (kind gift from Scott Gradia; Addgene plasmid # 55219)^19^ in frame with an N-terminal 6xHis tag followed by a TEV cleavage site, followed by generation of baculovirus.

The integrity of all plasmids was confirmed by Sanger sequencing.

### Immunofluorescence experiments

For immunofluorescence experiments, 143B, KO cells and derived FASTKD5-overexpressing cell lines were grown on coverslips for 24 hours. Cells were then fixed using 4% formaldehyde in PBS for 20 min at 37°C. Coverslips were washed 2 times with PBS, cells were permeabilized in 0.5% Triton in PBS for 15 min at room temperature and washed five times with PBS and blocked in PBS containing 5% BSA for a minimum of 10 minutes. Coverslips were then incubated with the indicated primary antibodies (**Table S1**) for one hour at room temperature, washed 3 times with PBS and incubated with the appropriate anti-species secondary antibodies coupled with Alexa fluorochromes (Thermo Fisher Scientific, 1:2000) and DAPI (Sigma, 1:2000) for 30 min at room temperature. Coverslips were washed three times with PBS and mounted with Fluromount-G (Thermo Fisher Scientific). Cells were imaged on an Olympus IX83 microscope connected with Yokogawa CSU-X confocal scanning unit, using UPLANSAPO 100x/1.40 Oil objective (Olympus) and Andor Neo sCMOS camera. Images were processed in Fiji ^20^.

### Whole cell extracts and mitoplast preparation

For SDS-PAGE, pelleted 143B, KO and over-expressing cells were resuspended in 1.5% n-dodecyl β-D-maltoside/PBS for 40 min on ice, centrifuged at 20000 g for 20 min at 4°C and protein concentration in supernatants was determined by Bradford assay. 2x Laemmli sample buffer was added to supernatants and samples were denatured at room temperature over-night or at 50°C for 10 min.

For Blue-Native PAGE (BN-PAGE), mitoplasts were prepared as described before ^21^. Briefly, pelleted 143B, KO and over-expressing cells were resuspended in PBS at a final concentration of 5 mg/ml. Digitonin was added to the cells at a ratio of 0.8 mg digitonin/mg of protein, and the samples were incubated on ice for 5 min, diluted to 1.5 ml with PBS, and centrifuged for 10 min at 10000 g, at 4°C. The pellet was resuspended in a buffer containing 75 mM Bis-Tris (pH 7.0), 1.75 M aminocaproic acid, and 2 mM EDTA at a final protein concentration of 2 mg/ml. 1/10 volume of 10% n-dodecyl β-D-maltoside was added to the samples, and the samples were incubated on ice for 20 min, followed by centrifugation at 20000 g for 20 min, at 4°C. Protein concentration in supernatants was determined by Bradford assay.

### Denaturing and native PAGE, immunoblot analysis

SDS-PAGE was used to separate whole cell extracts. 20 µg-50 µg of protein was run on either 10% or 12.5% polyacrylamide gels. BN-PAGE was used to separate individual OXPHOS complexes. 20 µg of mitoplasts were run on native 6-15% polyacrylamide gradient gels as previously described ^22^. Separated proteins were transferred to a nitrocellulose membrane and immunoblot analysis was performed with indicated antibodies (**Table S1**).

### Mitochondrial translation assay

Pulse-labeling of mitochondrial translation products in 143B, KO and over-expressing cells was performed with 100 μCi/ml of a [^35^S]-methionine/cysteine mix (Revvity), in DMEM lacking methionine and cysteine, and containing 100 μg/ml of a cytoplasmic translation inhibitor emetine, for 60 minutes as described in detail elsewhere ^23^. Cells were collected, resuspended in PBS to measure protein concentration using either Bradford or BCA assay. Total cellular protein (50 μg) was resuspended in loading buffer containing 93 mM Tris-HCl, pH 6.7, 7.5% glycerol, 1% SDS, 0.25 mg bromophenol blue/ml and 3% mercaptoethanol, sonicated for 3–8 s, loaded and run on 15-20% polyacrylamide gradient gels. After the run, the top part of the gel (above 70 kDa marker) was cut and stained with Coomassie Brilliant Blue R-250, to determine the protein loading. The rest of the gel was dried, and the labeled mitochondrial translation products were detected by direct autoradiography.

### RNA isolation, Northern-blot analysis and RT-qPCR

Total RNA was isolated from 143B, KO and over-expressing cells using the RNeasy Plus Kit (Qiagen) and extensively treated with DNase I to remove mtDNA. 5 μg of total RNA were separated on a denaturing MOPS/formaldehyde agarose gel and transferred to a nylon membrane. 300-500 bp long PCR products of individual mitochondrial genes were labeled with [α-^32^P]-dCTP (Revvity) using a Random Primed DNA labeling kit (Sigma). Hybridization was performed according to the manufacturer’s manual using ExpressHyb Hybridization Solution (Takara Bio) and the radioactive signal was detected using a Phosphorimager system. RT-qPCR analysis for all mitochondrial transcripts to determine the total mtRNA levels was performed at the Institute for Research in Immunology and Cancer (IRIC).

### DNA isolation and qPCR

Total DNA was isolated from 143B and KO cells using a DNeasy Blood and Tissue Kit (Qiagen). qPCR analysis was performed to determine mtDNA levels using four different qPCR assays (ND4, RNR2, D-loop, cyt b) at IRIC.

### Enzyme activity measurements

Spectrophotometric assays of whole cell extracts were used to measure enzyme activities as described previously ^24, 25^. Briefly, cells were grown on 10 cm plates to about 90% confluency, washed twice with PBS and scraped in 1 ml ice-cold PBS. Cells were collected by centrifugation at 10000 g for 1 min at 4°C. Cell pellets were resuspended in 50 mM triethanolamine pH 7.4, 1 mM EDTA and homogenized with motorized pestle for 15s. Citrate synthase activity was measured as an increase in absorbance at 412 nm at 30°C in 100mM Tris-HCl pH7.4, 25 µM acetyl-coenzyme A, 200 µM Ellman’s reagent (DTNB), 0.1% Triton X-100 with or without 70 µM oxaloacetic acid (pH 7.4) as a substrate. Cytochrome *c* oxidase (COX) activity was measured as a decrease in absorbance at 550 nm at 30°C in 50 mM potassium phosphate buffer (pH 7.0), 0.1% n-dodecyl β-D-maltoside and 50 µM reduced cytochrome *c* as a substrate. COX activity was normalized to citrate synthase activity.

### Cell growth assay

On day one 10000 cells were seeded in six-well culture dishes either in medium with or without uridine. After 1, 2, 3 and 4 days of culture, cells were trypsinized, homogenized, and counted using a Bio-Rad TC10 automated cell counter.

### Protein modelling

The Alphafold ^26^ protein prediction for FASTKD5 (AF-Q7L8L6-F1) was used to model individual point mutation using PyMol software (The PyMOL Molecular Graphics System, Version 3.0 Schrödinger, LLC).

### Recombinant FASTKD5 protein preparation

FASTKD5 lacking the predicted MTS (Δ1-27) was expressed in an insect cell expression system. Baculoviruses for protein expression in insect cells were generated in Sf9 and Sf21 cells. Large-scale expression of FASTKD5 was done in Hi5 insect cells.

FASTKD5 was expressed in an insect cell expression system using Hi-Five cells as previously described ^27, 28^. All protein purification steps were carried out at 4°C. Cells were harvested by centrifugation at 238 g for 30 minutes and resuspended in lysis buffer at pH 8.5 containing 50 mM Tris, 400 mM NaCl, 20 mM imidazole, 2 mM dithiothreitol (DTT), 10% glycerol and 1x protease inhibitor cocktail (Roche). Cells were lysed by sonification and subsequently centrifuged first in an SS34 rotor (Sorvall) at 47800 g for 30 minutes and the supernatant was then ultracentrifuged in a type 45 Ti Fixed-Angle rotor for 1 hour at 235 000 g. The supernatant was filtered through 0.8 μm filter and applied to a HisTrap HP 5 ml column (Cytiva) equilibrated with lysis buffer. The column was washed with 5 column volumes (CV) of high-salt wash buffer (50 mM Tris, 1 M NaCl, 30 mM imidazole, 10% glycerol, 2 mM DTT, pH 8.5), followed by 5 CV of lysis buffer. Bound proteins were eluted with elution buffer (50 mM Tris, 400 mM NaCl, 300 mM imidazole, 10% glycerol, 2 mM DTT, pH 8.5) over 15 column volumes (CV). 2 mg of recombinant 6xHis-TEV protease (in house) was added, and the eluate was dialyzed overnight against 50 mM Tris, 300 mM NaCl, 10% glycerol, 2 mM DTT, pH 8.5 using a 7-kDa molecular weight cut-off (MWCO) SnakeSkin dialysis tubing (Thermo Fisher Scientific). To remove uncleaved protein, the imidazole concentration was adjusted to 30 mM and the eluate was reapplied to the HisTrap HP 5 ml column equilibrated with buffer containing 50 mM Tris, 300 mM NaCl, 30 mM imidazole, 10% glycerol, 2 mM DTT, pH 8.5 and washed with 5 CV of the same buffer. The flow-through and wash were collected and diluted with a low-salt buffer (50 mM Tris, 20 mM NaCl, 10% glycerol, 2 mM DTT, pH 8.5) to a final salt concentration of 150 mM. The sample was applied to a HiTrap Heparin 5 mL column (Cytiva) equilibrated with 50 mM Tris, 150 mM NaCl, 10% glycerol and 2 mM DTT, pH 8.5 and washed with 5 CV of the same buffer. Bound proteins were eluted with a gradient of 150 mM to 1000 mM NaCl in 50 mM Tris, 10% glycerol, 2 mM DTT, pH 8.5. Fractions containing FASTKD5 according to SDS-PAGE were pooled, concentrated using Amicon 30 kDa MWCO Centrifugal filters (Sigma) and further purified using a Superdex 200 Increase 10/300 GL column (Cytiva) equilibrated with 20 mM Tris, 350 mM NaCl, 5% glycerol, 5 mM DTT, pH 8.5. Fractions containing FASTKD5 were concentrated using Amicon Ultra 0.5 ml 30K Centrifugal Filters (Sigma), aliquoted, flash-frozen in liquid nitrogen and stored at −80 °C.

### *In vitro* RNA processing assay

The cleavage assays were carried out in 10 μl reactions containing RNA cleavage assay buffer (100 mM Tris-HCl pH 7.5, 100 mM NaCl, 4 mM DTT), 200 nM 3’-end Cy3 labeled RNAs (**Table S1**) and 0.3 µM FASTKD5. For the cleavage assays, the RNA cleavage assay buffer and FASTKD5 protein were first mixed and incubated at 30°C for 5 min. RNA was added, and reactions were incubated at 30°C for a further 60 min. The reactions were quenched with an equal volume of 2× RNA loading buffer (6.4 M urea, 2× Tris-borate-EDTA buffer, 50 mM EDTA pH 8.0) and 1.6 U of recombinant proteinase K (New England Biolabs) was added. The reactions were incubated at 30°C for 60 min or at 50°C for 10 min and stored at −20°C until analysis by urea–PAGE. Before urea–PAGE, the reactions were thawed and incubated at 95°C for 3 min. A 10 μl portion of each reaction was loaded onto 7 M urea, 10% polyacrylamide (bis-acrylamide to acrylamide ratio 1:19) denaturing gels in 1× Tris-borate-EDTA buffer. After electrophoresis, gels were imaged using an INTAS imaging system (INTAS Science Imaging, Germany). microRNA marker (NEB) and Low Range ssRNA ladder (NEB) were used to estimate the size of the RNA bands; Cy3-adds approximately 3 nt to the size of the actual RNA.

### General statistical analysis

Unless indicated otherwise in the figure legends, all experiments were performed at least in biological triplicates, and results are presented as mean ± standard deviation (SD) or standard error of the mean (SEM) of absolute values or percentages of control. For the comparison of multiple groups, two-way ANOVA with a Dunnett correction for multiple comparisons was performed. A p < 0.05 was considered significant. The exact p values are indicated in the graphs, except for the cell growth assay, where p values are as follows: * p < 0.05, ** p < 0.01, *** p < 0.001, and **** p < 0.0001.

## Results

### FASTKD5 KO cells completely lack processed non-canonical transcripts

Previous investigations showed that FASTKD5 was required for the complete processing of non-canonical pre-mRNAs (not flanked by tRNAs) in the primary mitochondrial polycistronic transcript ^8, 9^. To investigate the precise molecular basis for FASTKD5 activity we first sought to create and characterize a cell line null for FASTKD5 which we could then use as an assay system to assess the functionality of specific domains/amino acid residues in the FASTKD5 protein. We used CRISPR/Cas9 editing to create a knockout (KO) line in 143B cells and propagated them in medium containing pyruvate and uridine, which supports the replication of cells with deficient OXPHOS activity. We isolated and characterized two independent KO clones. Immunoblot analysis of the steady-state levels of mtDNA-encoded polypeptides showed specific decreases in COX I and cyt b, both encoded in non-canonical pre-mRNAs, but not ND1 or ATP6 (Fig. 1A). Retroviral expression of a FLAG-tagged wild-type FASTKD5 construct rescued this phenotype, demonstrating the specificity of the CRISPR/Cas9 editing. Immunofluorescence analysis using an antibody specific for COX I, showed complete loss of COX I at the single cell level in the FASTKD5 KO cells (Supp Fig. S1B) and rescue of the phenotype only in those cells expressing the FLAG-tagged wild-type construct (Fig. 1G).

**Figure 1.**
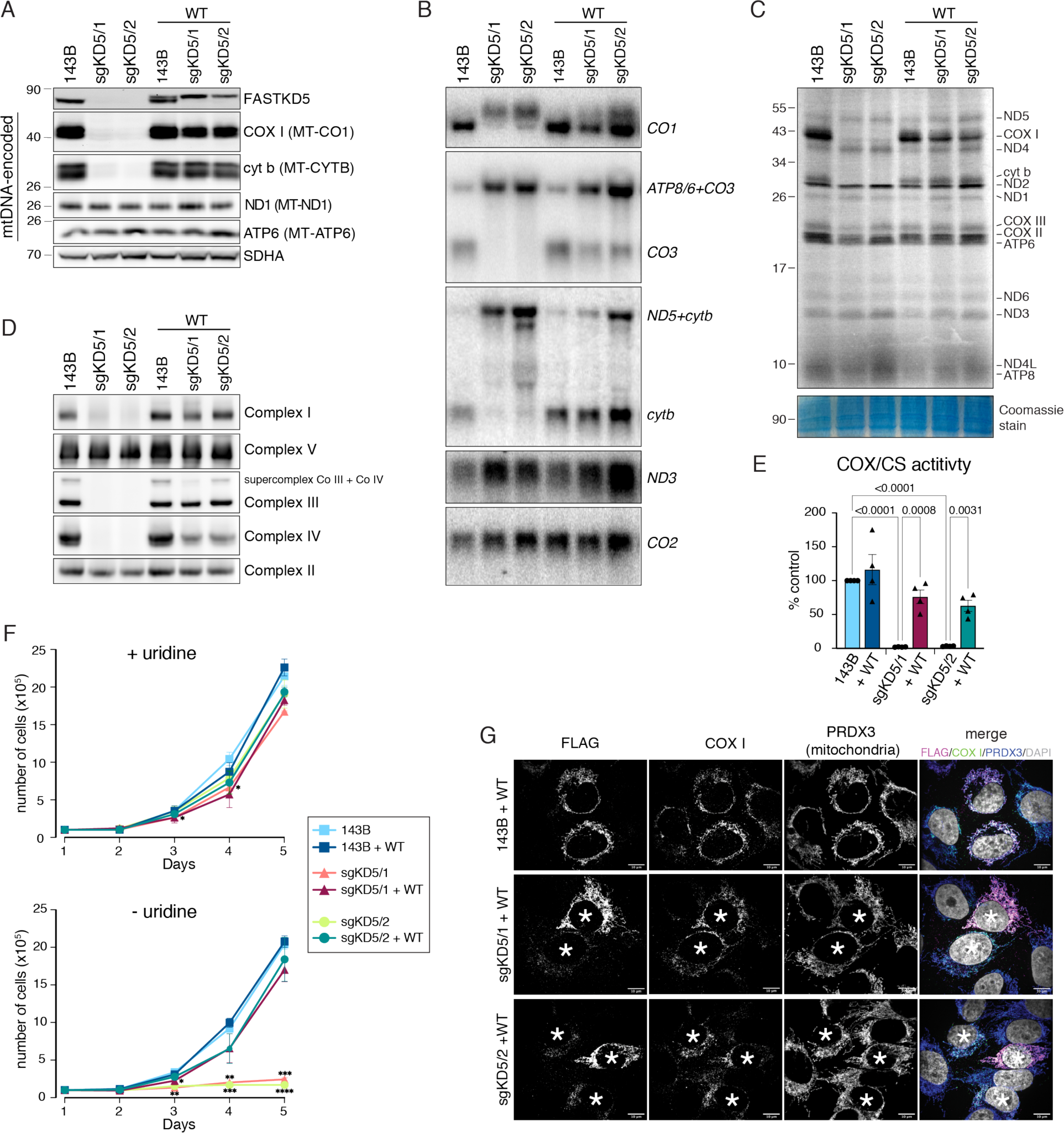
Lack of FASTKD5 protein leads to a loss of mtRNA processing at non-canonical sites, and a lack of synthesis of COX I and cyt b resulting in a combined OXPHOS deficiency; phenotypes that are rescued by re-expression of wild-type FASTKD5. (**A**) Steady-state levels of FASTKD5 protein and selected proteins encoded by mtDNA assessed by immunoblotting in 143B cells, two KO clones (sgKD5/1 and sgKD5/2) and same cells over-expressing wild-type (WT) FASTKD5-3xFLAG. The names in parenthesis indicate the gene names. SDHA was used as a loading control. Molecular weight markers (in kDa) are indicated on the left. (**B**) Defective processing of mitochondrial transcripts was evaluated by Northern blot analysis in 143B cells, two KO clones (sgKD5/1 and sgKD5/2) and same cells over-expressing wild-type (WT) FASTKD5-3xFLAG. The canonical transcripts (*ND3, CO2*) are used as loading control. (**C**) Impaired synthesis of COX I and cyt b polypeptides in FASTKD5 KO clones (sgKD5/1 and sgKD5/2) following ^35^S-methionine/cysteine incorporation into newly synthesized mitochondrial polypeptides in the presence of emetine to inhibit cytoplasmic protein synthesis. A Coomassie total protein staining served as a loading control. Molecular weight markers (in kDa) are indicated on the left. (**D**) OXPHOS assembly defect in FASTKD5 KO clones (sgKD5/1 and sgKD5/2) was assessed by BN-PAGE analysis with complex specific antibodies. Complex II was used as a loading control. (**E**) Lack of cytochrome *c* oxidase (COX) activity in FASTKD5 KO clones (sgKD5/1 and sgKD5/2) determined by spectrophotometric enzymatic assay, shown as a COX/citrate synthase (COX/CS) ratio and normalized to the COX/CS ratio in 143B cells, which was set to 100%. Two-way ANOVA with a Dunnett correction for multiple comparisons was performed, and significant p-values are indicated. (**F**) The growth of FASTKD5 KO clones (sgKD5/1 and sgKD5/2) is dependent on uridine, as measured by cell growth over 5-day period. Two-way ANOVA with a Dunnett correction for multiple comparisons was performed, and significant p-values are indicated; *p < 0.05, **p < 0.01, ***p < 0.001, ****p < 0.0001 (**G**) Rescue of COX I expression assessed at the single-cell level by immunofluorescence in 143B cells and two KO clones (sgKD5/1 and sgKD5/2) over-expressing wild-type (WT) FASTKD5-3xFLAG. Cells expressing WT-FASTKD5 were detected by anti-FLAG antibody (marked with asterisks), PRDX3 was used as a mitochondrial control.

We next proceeded to investigate the effects of the FASTKD5 KO on RNA processing by Northern blot analysis (Fig. 1B) which showed the near complete loss of the mature, processed forms of the *CO1*, *CO3* and *cytb* mRNAs, a phenotype rescued by expression of the FLAG-tagged wild-type construct. The lack of processing resulted in a decrease of total mRNA levels (processed and unprocessed) for the *CO1* transcript as determined by RT-qPCR analyses (Supp Fig. 1C), but not any other transcript. We conclude that the complete loss of FASTKD5 function specifically affects the processing of only the non-canonical pre-mRNAs in the primary polycistronic transcript.

We subsequently examined the effects of the FASTKD5 KO on the translation of mtDNA-encoded polypeptides by pulse-labeling with [^35^S]-Met/Cys in the presence of emetine, an inhibitor of cytoplasmic translation (Fig. 1C). Quantification of these results (Supp Fig. S1F) showed that the translation of COX I and cyt b was specifically decreased, to near zero, consistent with the results of the Northern blot analysis (Fig. 1B) demonstrating that the presence of 5’-UTRs on these transcripts prevents translation on mitochondrial ribosomes. The translation of COX III was, however, not affected in the KO cells, although the level of the mature *CO3* mRNA was virtually undetectable by Northern blot. This directly demonstrates that a tricistronic mRNA (*ATP8/6+CO3*) can be translated by the mitochondrial ribosome as has been suggested previously^29^.

Analysis of the assembly of the OXPHOS complexes by BN-PAGE demonstrated that complexes III and IV failed to assemble as expected from the mitochondrial translation experiments (Fig. 1D). However, we also observed a near complete lack of fully assembled complex I even though the mtDNA-encoded subunits of this complex were translated at near normal levels. The complex I assembly defect can be ascribed to the complete absence of complex III, whose presence is necessary for the biogenesis/stability of complex I ^30, 31^. The maximal enzyme activity of Complex IV (cytochrome *c* oxidase) was also reduced as expected from the BN-PAGE results (Fig. 1E).

To investigate the impact of the deletion of FASTKD5 on the steady-state levels of other proteins involved in posttranscriptional regulation of mitochondrial gene expression, we performed immunoblot analyses for a wide spectrum of proteins that localize to the mitochondrial RNA granule^8^ and are involved in RNA processing, the pseudouridine synthase module^32^ as well as proteins involved in mitochondrial transcription and translation (Supp Fig. S1A). None of these proteins were altered by deletion of FASTKD5 or by re-expression of the FLAG-tagged construct. The levels of mtDNA were also not significantly altered by the FASTKD5 KO (Supp Fig. S1E), and neither was the appearance or number of mitochondrial nucleoids and RNA granules (Supp Fig. S1D).

To test whether the OXPHOS assembly defects in the FASTKD5 KO cells made the cells uridine auxotrophs, we omitted uridine from the medium and observed that the growth rate was severely depressed, an effect that was rescued by expression of a wild-type FASTKD5 construct (Fig. 1F).

### Functional domains and amino acid residues of FASTKD5

We suspected that all domains in FASTKD5 would be necessary for function; however, we formally investigated this by deleting them individually and testing whether any of the deletion mutants would restore function in FASTKD5 KO cells. Deletion of any of the domains (MTS, heptatricopeptide repeats, FAST1, FAST2, and RAP domains) resulted in loss of function as analyzed by immunoblot analysis or immunofluorescence at the single cell level (Supp Fig. 2A-D).

**Figure 2.**
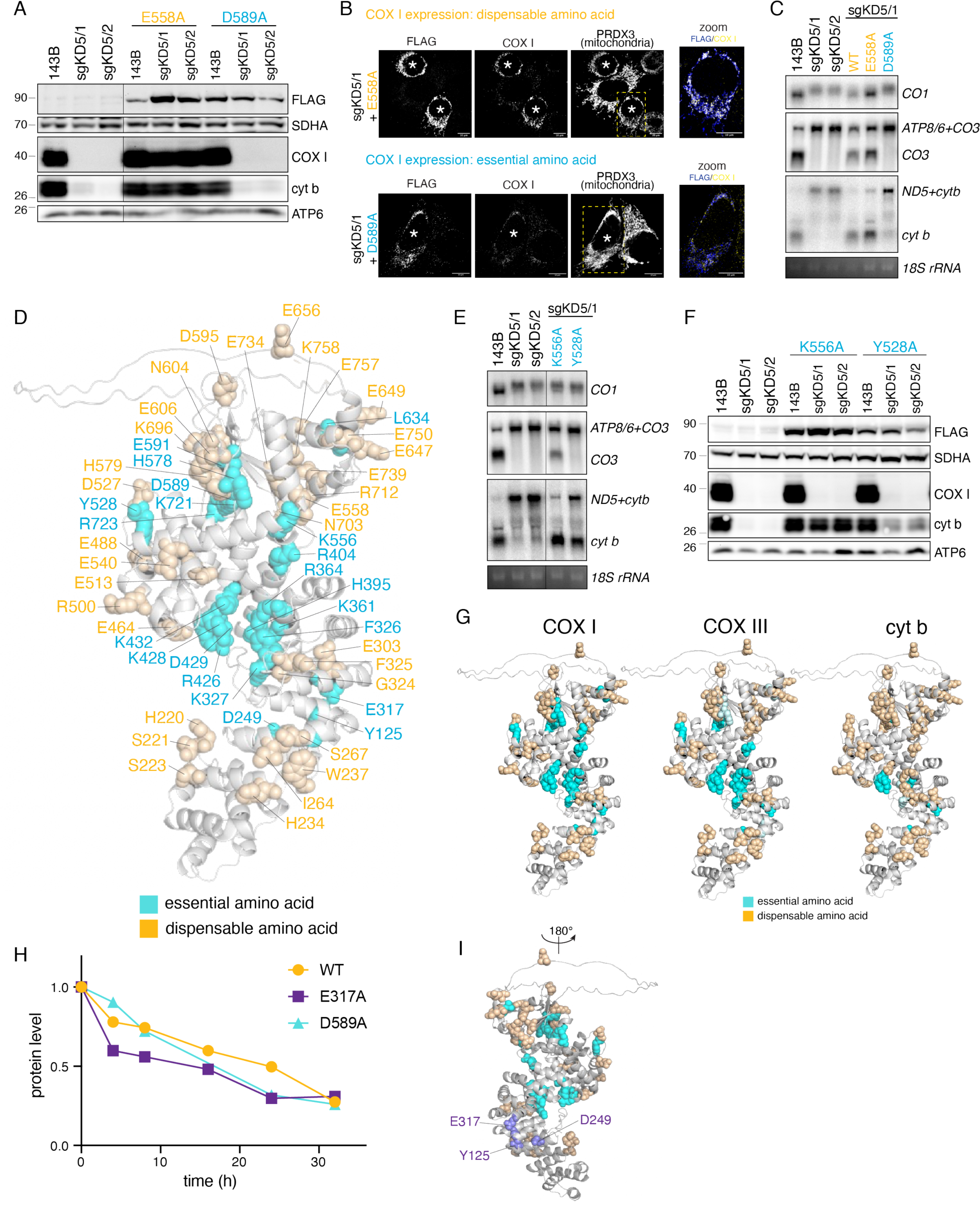
Single amino acid mutational analysis identifies 21 amino acids essential for FASTKD5 function in mtRNA processing. (**A**) Immunoblot analysis of 143B cells and two KO clones (sgKD5/1 and sgKD5/2) reconstituted with either E558A (dispensable amino acid) or D589A (essential amino acid) FASTKD5-3xFLAG protein variant. SDHA was used as a loading control. Molecular weight markers (in kDa) are indicated on the left. (**B**) Rescue of COX I expression (for E558A variant) or lack of rescue for the D589A variant assessed at the single-cell level by immunofluorescence in a KO clone (sgKD5/1) reconstituted with the indicated FASTKD5-3xFLAG protein variant detected by the anti-FLAG antibody (marked with asterisks). PRDX3 was used as a mitochondrial control. Right panel, zoom of the indicated area showing the merge of the FLAG/COX I staining. (**C**) mtRNA processing is rescued in E558A variant, but not in D589A variant. Processing of non-canonical transcripts determined by Northern blot analysis in the indicated cells. UV-stain of 18S rRNA was used as a control. (**D**) Modelling of essential (in blue) and dispensable (in orange) amino acids for COX I expression on the Alphafold-predicted FASTKD5 structure. For clarity, the unstructured sequence of 1-100 amino acids was omitted from the illustration. (**E**) Variability in *ATP8/6+CO3* and *ND5+cytb* processing in cells reconstituted with essential amino acids for *CO1* processing (K556A, Y528A) determined by Northern blot analysis. UV-stain of 18S rRNA was used as a control. (**F**) Rescue of cyt b steady state levels in cells reconstituted with essential amino acids for COX I expression (K556A, Y528A). SDHA was used as a loading control. Molecular weight markers (in kDa) are indicated on the left. (**G**) Modelling of essential (in blue) and dispensable (in orange) amino acids for COX I, COX III or cyt b expression on the Alphafold-predicted FASTKD5 structure (same as in **D**). The residues in light blue provided only partial rescue in the KO cells, and we scored them as essential amino acids. (**H**) The E317A-FASTKD5 variant is less stable than the WT-FASTKD5 protein. Protein levels of WT, D589A and E317A variants were determined by immunoblotting, after inhibition of cytosolic translation for the indicated time, and normalized to actin levels and further normalized to the protein level at time zero, which was set to 1. (**I**) Modelling of amino acids responsible for stability of FASTKD5 protein (indicated in purple) on the Alphafold-predicted FASTKD5 structure (same as in **D** rotated vertically by 180°).

We hypothesized that the heptatricopeptide repeats likely contain RNA-binding/recognition properties and that the region containing the endonuclease-like fold may contain the active site. We constructed a large series of mutants in which we mutated single amino acid residues using as a guide those that are evolutionarily conserved in FASTKD5 proteins from other species and in other members of the FASTKD family, as well as several variants of unknown significance identified in probable FASTKD5 patients. We made a total of 53 amino acid substitutions, mutating the conserved residues mostly to alanine (Table S1) and tested all of them for rescue of FASTKD5 function by immunoblotting and immunofluorescence with an anti-COX I antibody (Fig. 2A, B, Supp Fig. S3). Fig 2. A, B shows typical results from a dispensable and an essential amino acid residue. We confirmed that the residues we mapped as essential all affected pre-mRNA *CO1* processing (Fig. 2C). We modeled all the profiled variants on the Alphafold predicted model of FASTKD5 (Fig. 2D), creating a map of 21 essential and 32 dispensable amino acids for the expression of the COX I polypeptide. Northern blot analysis of processing of *ATP8/6+CO3* and *ND5+cytb* transcripts surprisingly showed that some amino acid residues that were essential for processing of *CO1* were dispensable for processing of *CO3*, and further, some residues essential for processing of the latter were dispensable for processing of *cytb* (Fig. 2E, Supp Fig. S4A). Rescue of *cytb* processing resulted in the rescue of cyt b synthesis (Supp Fig. S4B) and cyt b steady state levels (Fig. 2F, Supp Fig. S3). Maps of the essential vs. dispensable residues for the three different processing events are shown on Alphafold-predicted structures (Fig 2G). Remarkably, only seven of those sites necessary to process *CO1* were necessary to process *cytb*, and no dispensable residues for *CO1* processing appeared as essential for *cytb* processing. Immunoblot analysis of the expression of individual variants identified three variants that were expressed significantly less than the WT FASTDK5 or the other variants, suggesting that they are important for protein stability. To formally address this, we measured the half-life of one of these variants (E317A) and compared it to WT protein and the D589A variant (Fig. 2H) following inhibition of cytosolic translation with cyclohexidime. The half-life of E317A variant was ∼5 hours, compared to 17 hours for D589A and 49 hours for WT protein. The half-life of endogenous FASTKD5 protein in 143B cells was 43.7 hours. We conclude that there are several variants critical for the stability of FASTKD5 protein (Fig. 2I) and there is a hierarchical dependence of FASTKD5 function for processing non-canonical pre-mRNAs that is context (pre-mRNA) dependent.

### FASTKD5 accurately processes non-canonical pre-mRNAs *in vitro*

To test whether FASTKD5 on its own can accurately process pre-mRNA substrates we purified FASTKD5 protein (lacking the MTS) from insect cells and tested whether we could obtain specific cleavage on synthetic RNA substrates. To this end we used synthesized single stranded RNAs, that were either potential substrates or downstream sequences, tagged at the 3’end with the fluorophore Cy3. FASTKD5 processed all three substrates at the expected sites, producing a band running at 27 nt (Fig. 3A). For the pre-mRNA substrates for *ATP8/6+CO3* and *ND5+cytb*, several other smaller cleavage products were also observed (Fig. 3A). The quantity of processed RNA was FASTKD5 dose-dependent (Fig. 3B). We conclude that FASTKD5, on its own, possesses an activity that can process client RNA substrates appropriately. The fact that it does not process RNA sequences that are not 5’ UTRs shows that this is not due to non-specific nuclease activity; however, the observation that additional cleavage products are present other than those expected on authentic substrates, suggests that other factors may be necessary for specificity or efficient cleavage.

**Figure 3.**
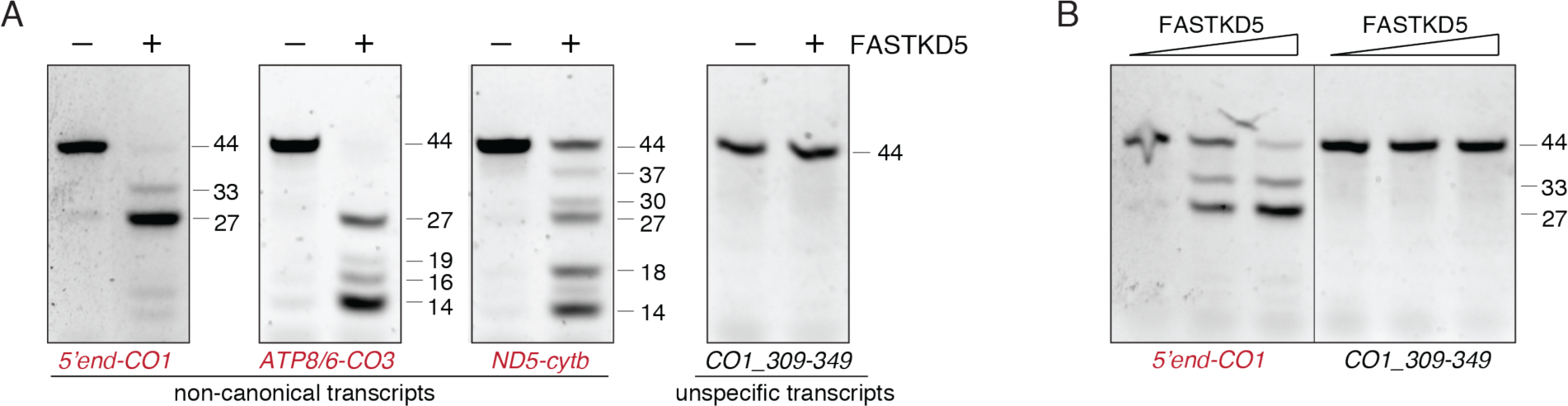
FASTKD5 cleaves non-canonical pre-mRNA transcripts in an *in vitro* cleavage assay. (**A**) Recombinant FASTKD5 protein was incubated with indicated 3’-end Cy3-labelled RNAs, and the resulting products were separated on 10% 7 M urea-denaturing PAGE. Estimated sizes of individual RNAs (nt) based on the migration of RNA markers are indicated on the right. Non-cleaved RNA runs at 44 nt (Cy3-adds approximately 3 nt to the size of the actual RNA). The expected cleaved RNA runs at 27 nt (the predicted RNA product is 24 nt) (**B**) Processing of 5’*-*end *CO1* transcript is dependent on the FASTKD5 protein levels. Estimated sizes of individual RNAs (nt) based on the migration of RNA markers are indicated on the right.

## Discussion

This study demonstrates that a knockout of FASTKD5 results in a severe deficiency in the assembly of multiple OXPHOS complexes resulting from an inability to process the non-canonical pre-mRNAs in the primary polycistronic mitochondrial transcript, extending our previous observations^8^. This prevents translation of at least two mRNAs, encoding *CO1* and *cytb*, supporting observations that mRNAs with 5’-UTRs cannot generally load efficiently onto mitoribosomes to form an initiation complex ^29, 33^. Aside from the two known bicistronic transcripts (*ATP8/6* and *ND4L/ND4*), one exception to this general rule is the *CO3* mRNA which can load on to mitochondrial ribosomes as a tricistronic *ATP8/6/CO3* transcript^29^. Despite undetectable levels of the mature, processed *CO3* mRNA in the FASTKD5 KO cells, *CO3* mRNA is translated normally downstream of the *ATP8/6* overlapping reading frames. The total steady-state levels of *CO1* and *cytb* were the only mRNAs in the FASTKD5 cells that were significantly reduced likely reflecting increased turnover due to their inability to associate with the mitoribosome, and the pulse translation experiment demonstrated that these were the only mRNAs whose synthesis was significantly reduced (to undetectable levels) in the KO cells. These processing defects resulted in a complete failure to assemble Complexes III and IV, and because the stability of Complex I depends on the assembly of Complex III,^30, 31^ a complete loss of fully assembled Complex I. This loss of OXPHOS capacity rendered the KO cells uridine auxotrophs as the synthesis of pyrimidines is intimately coupled to a functional OXPHOS system.

A previous study of FASTKD5 function in a haploid cell line (Hap1) in which the authors used CRISPR/Cas9 editing to disrupt FASTKD5, did not show a defect in the translation of *cytb* or in the assembly of Complexes III or I^9^, and while this may partly reflect cell specific responses to loss of FASTKD5 function, it is clear from their data that their KO cells still harboured residual FASTKD5 protein. Similarly, in our previous study in 143B cells, in which FASTKD5 protein levels were decreased by siRNA-mediated knock-down ^8^ processing of *ND5/cytb* was not affected, implying that even a small amount of FASTKD5 protein may be sufficient to correctly process *cytb* pre-mRNA.

Whether FASTKD5 possesses endonuclease activity or whether it recognizes its substrates and recruits other proteins with such activities to mature the non-canonical substrates has been the subject of some speculation ^8–10^. We clearly demonstrate, using an *in vitro* reconstituted system, that FASTKD5 possesses an activity sufficient to correctly process three different synthetic pre-mRNA substrates of the primary polycistronic mitochondrial transcript. Thus, complete maturation of the primary mitochondrial transcript requires the concerted activity of three different RNA processing factors: RNase P, RNase Z and FASTKD5. Unlike pre-mRNAs that are flanked by tRNAs that assume a characteristic tertiary structure, there does not appear to be any common sequence or RNA structure that might denote the processing sites for FASTKD5. Both RNase P and RNase Z utilize two additional subunits (TRMT10C, SDR5C1) as RNA recognition platforms to correctly process their substrates ^3, 5–7^. While PRORP can process some pre-mRNAs without these additional factors, the processing rate is 10 times slower and not all substrates can be processed ^34^. We used an endpoint assay to assess FASTKD5 activity and so it is not possible to estimate turnover. The idea that RNA processing occurs co-transcriptionally would seem to imply that all primary processing events have similar kinetics. Recent estimates from the Churchman lab ^29^ indicate that the rate of transcription of the polycistronic transcript is <1kb/min and that the half-lives of the pre-mRNAs (canonical and non-canonical) are in the range of 1-39 min consistent with the idea that the two processes are coupled. Bhatta et al.^7^ showed experimentally that the order of tRNA processing proceeds with 5’ processing by RNase P followed by 3’ cleavage by RNase Z. The fact that processing of pre-*CO3* and pre-*cytb* in our *in vitro* reconstituted system is not completely specific for single cleavage products strongly suggests that other factors might be involved in creating a platform for RNA recognition or be necessary for processivity. If in fact FASTKD5 uses a platform on which to process its substrates, then the nature of this may determine the order in which canonical and non-canonical cleavage events occur. An *in vivo* tissue-specific knockout of PRORP (MRPP3) showed that the non-canonical pre-mRNAs were not processed when 5’-tRNA cleavage was absent ^35^, suggesting that tRNA excision was necessary for FASTKD5 function, but whether this is a direct or indirect effect of the loss of canonical RNA processing remains unknown.

One other member of the FASTKD family, FASTKD4, has also been implicated in processing one non-canonical pre-mRNA, the *ND5/cytb* junction^9, 11^; however, FASTKD4 KO cells do not result in complete abrogation of this processing event and there is no observable deficiency in the synthesis of any of the mtDNA-encoded polypeptides in pulse translation experiments. Double KO (FASTKD4/5) Hap1 cells exhibit more severe defects in this processing event than do the single KO cells, but there is apparently no defect in the translation of either mRNA. It is clear from the data we present here that FASTKD5 is essential for the processing of the pre-mRNA *ND5/cytb* junction and that endogenous FASTKD4 cannot rescue this activity. Whether FASTKD4 somehow collaborates with FASTKD5 in this processing event will require further investigation.

A surprising outcome of our FASTKD5 mutational analysis is that while many amino acid residues tested are required for processing pre*-CO1*, a smaller number of the same essential residues are required to process pre*-CO3* and fewer still pre-*cytb*. In addition, several other cleavage products were observed for *CO3* and *cytb* in the *in vitro* assay. These data suggest that FASTKD5 cleavage of its substrates may not be one size fits all, and perhaps different substrates require or have recruited other proteins to assist in the cleavage or to confer specificity. It is likely that the heptatricopeptide repeat coiled-coil domains of FASTKD5 play a role in positioning the RNA substrate for cleavage and mutations in these abrogate function. To gain some insight into the nature of the endonuclease active site of FASTKD5, we tested all evolutionary conserved aspartate/glutamate residues. Our mutational analysis identified four aspartate/glutamate residues that were essential for processing of all three pre-mRNAs; however, they do not cluster in one domain of the protein (Supp Fig. S5A). As the essential positively charged residues (Fig. 2D, G) that are predicted to bind RNA cluster mostly near aspartate-429, we can speculate that this region contains the RNA binding/recognition site. As FASTKD1 and FASTKD4 were suggested to contain an endonuclease fold similar to the VSR endonuclease from *E. coli* ^11^ we modelled VSR onto the FASTKD5 protein structure predicted by Alphafold (Supp Fig. S5B). VSR aligned with the FAST2/RAP domain region of FASTKD5, which contains the D589 residue essential for processing all the synthetic RNA substrates we tested. While these models are suggestive, a solution to the structure of FASTKD5 will be necessary to definitively identify an active site for FASTKD5 and a mechanism for processing the non-canonical pre-mRNAs in the primary polycistronic transcript.

## Supporting information

Supplemental materials for FASTKD5

## Data and code availability

All reagents generated in this study (plasmids and cell lines) are available from the Lead Contact upon request. All Source Data is either included in the manuscript or will be provided upon request.

## Supplementary data

Figures S1-S5, and Table S1

## Acknowledgements

We thank Kathleen Daigneault for her technical assistance.

Author contributions: HA performed biochemical characterization, cell biology and *in vitro* studies. WW performed mitochondrial translation and Western blot assays. HA and EAS designed and led the project. AV and HSH prepared the recombinant protein and provided input into experimental plan. All authors contributed to data interpretation and read, edited, and approved the manuscript.

## Funding

This research was supported by a Canadian Institutes of Health Research grant # FRN178373 (to EAS), by the Deutsche Forschungsgemeinschaft under Germany’s Excellence Strategy EXC 2067/1-390729940 (to H.S.H), FOR2848 (P10 to H.S.H.), SFB1565 (Project number 469281184, P13 to H.S.H.) and by the European Union (ERC Starting Grant MitoRNA, grant agreement no. 101116869 to H.S.H.). A.V. is a doctoral student of the Ph.D. program “Molecular Biology” – International Max Planck Research School and the Göttingen Graduate School for Neurosciences, Biophysics, and Molecular Biosciences (GGNB) (DFG grant GSC 226) at Georg-August University of Göttingen. Views and opinions expressed are however those of the author(s) only and do not necessarily reflect those of the European Union or the European Research Council Executive Agency. Neither the European Union nor the granting authority can be held responsible for them.

## Conflict of interest statement

The authors declare no competing interests.

## Notes

### Competing Interest Statement

The authors have declared no competing interest.

