## Supplemental materials for FASTKD5 for "FASTKD5 processes mitochondrial pre-mRNAs at non-canonical cleavage sites"

**Supplementary Materials:**

**Supplementary Figures S1-S5**
**Supplementary Table S1**

Supplementary Figure S1

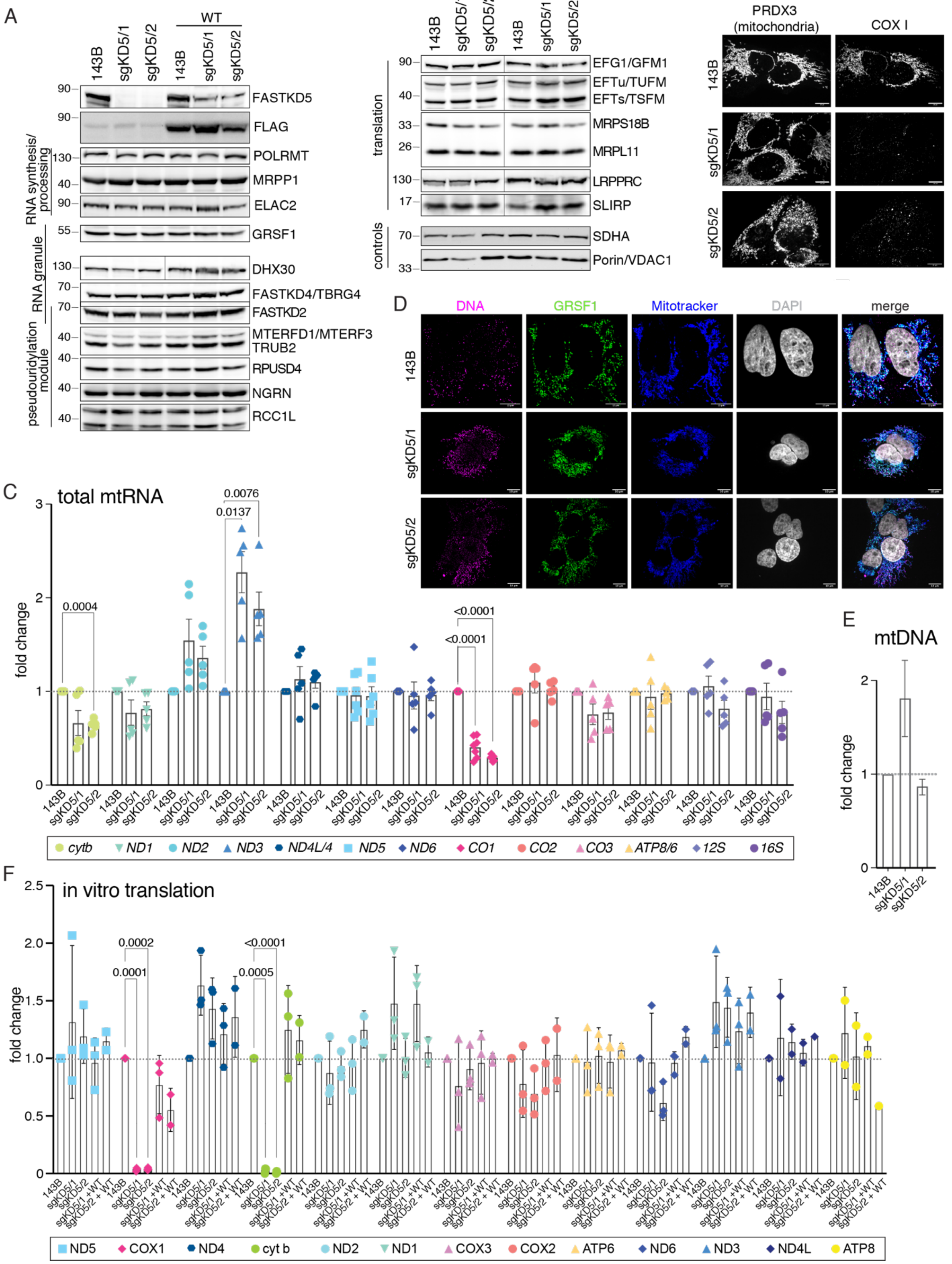

**Supplementary Figure S1** (related to **Figure 1**)

(**A**) Steady-state levels of FASTKD5 protein, FLAG and proteins involved in mitochondrial RNA metabolism or mitochondrial translation were assessed by immunoblotting in 143B cells, two KO clones (sgKD5/1 and sgKD5/2) and same cells rescued with a wild-type (WT) FASTKD5-3xFLAG. SDHA and VDAC1 were used as loading controls. Molecular weight markers (in kDa) are indicated on the left. (**B**) FASTKD5 KO clones (sgKD5/1 and sgKD5/2) show negligible levels of COX I expression on immunofluorescence. (**C**) RT-qPCR quantification of total mtRNA levels (processed + unprocessed) for all mitochondrial transcripts, as well as 12S (*RNR1*) and 16S (*RNR2*) rRNAs in 143B cells and two KO clones (sgKD5/1 and sgKD5/2). Two-way ANOVA with a Dunnett correction for multiple comparisons was performed (compared to 143B cells, which were set to 1), and significant p-values are indicated. (**D**) Nucleoids and RNA granules are normal in FASTKD5 KO cells as determined by immunofluorescence analysis using anti-DNA antibody (nucleoids) and anti-GRSF1 antibody (RNA granules). (**E**) mtDNA levels were measured by qPCR in 143B cells and two KO clones (sgKD5/1 and sgKD5/2) in duplicate. (**F**) Quantification of mitochondrial translation assay performed in biological triplicates (one example is shown in **Figure 1C**) normalized to Coomassie total protein staining. Two-way ANOVA with a Dunnett correction for multiple comparisons was performed (compared to 143B cells, which were set to 1), and significant p-values are indicated.

Supplementary Figure S2

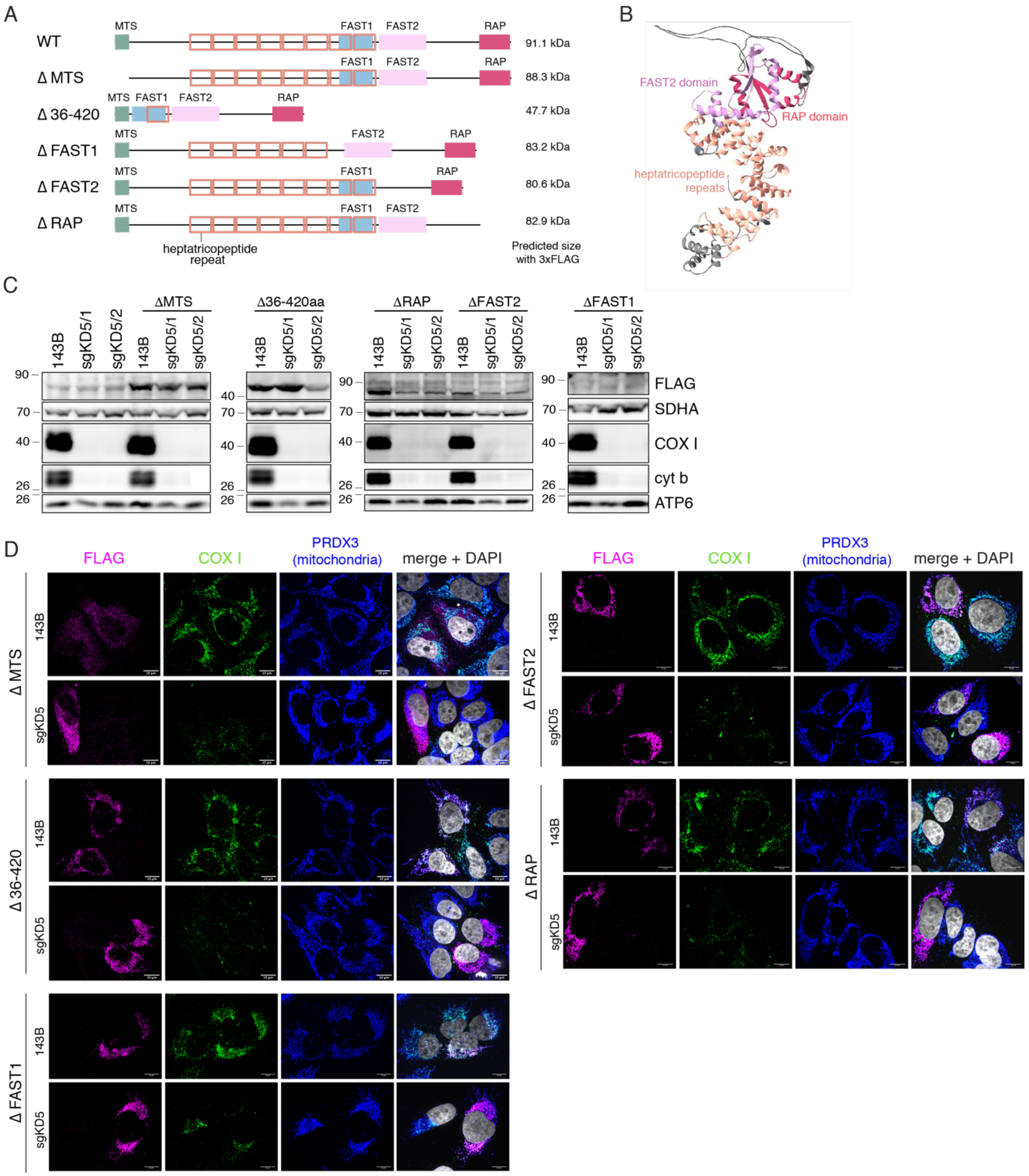

**Supplementary Figure S2** (related to **Figure 2**)

(A) Schematic representation of WT-FASTKD5 protein and individual deletion variants, indicating mitochondrial targeting sequence (MTS), previously described protein domains

(FAST1, FAST2 and RAP) and predicted heptatricopeptide repeats. Expected molecular weight of the deletion variants is indicated on the right. (**B**) Modelling of indicated domains on Alphafold-predicted FASTKD5 structure. For clarity, the unstructured sequence of 1-100 amino acids was omitted from the picture. (**C, D**) All domains are important for FASTKD5 function as reconstitution with the individual deletion variants did not rescue the expression of COX I or cyt b as assessed by Western blot analysis (**C**) or by immunofluorescence at the single cell level (**D**). SDHA was used as a loading control. PRDX3 was used as a mitochondrial marker. Molecular weight markers (in kDa) are indicated on the left (**C**).

Supplementary Figure S3

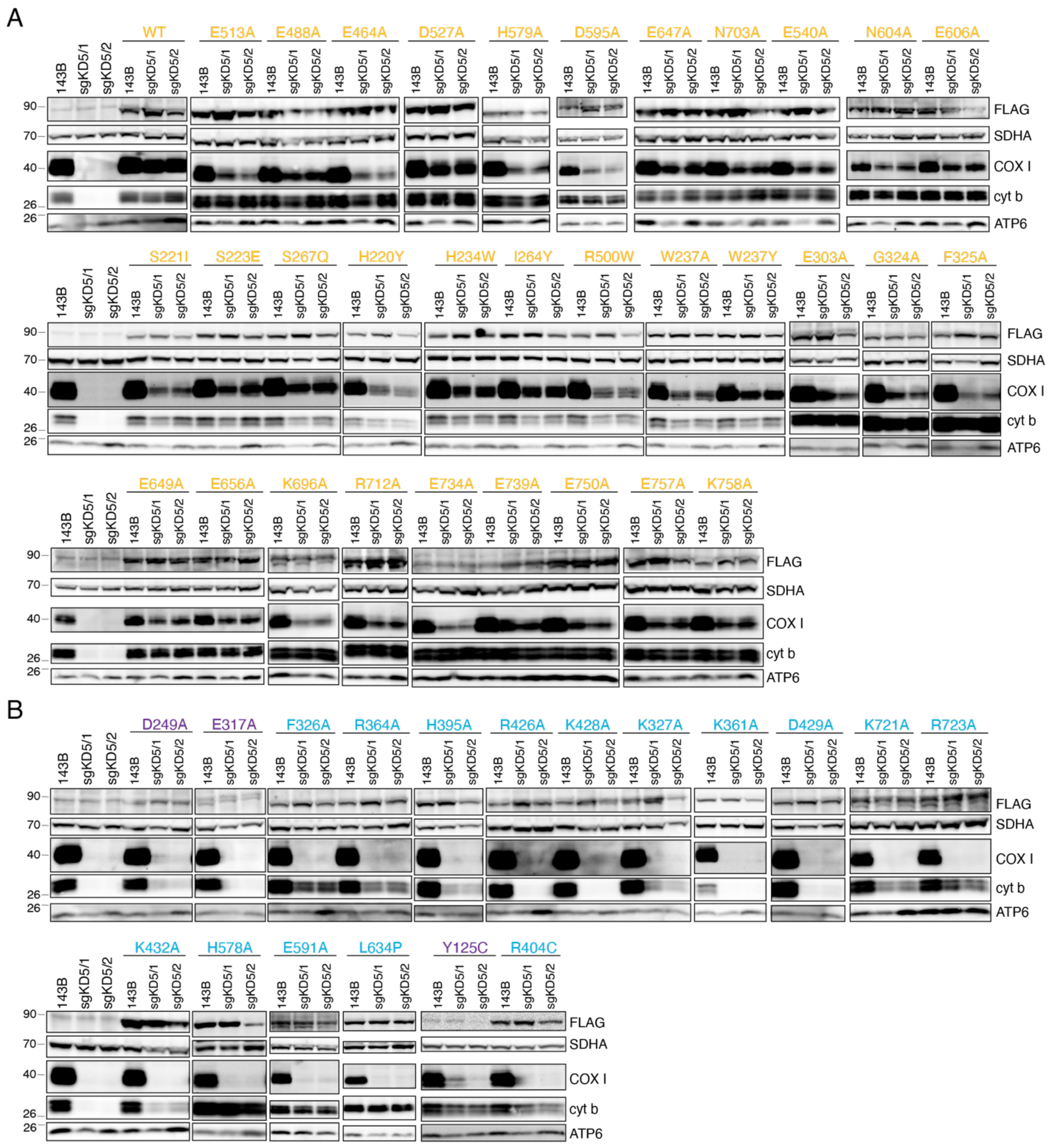

**Supplementary Figure S3** (related to **Figure 2**)

Western blot analysis of 143B cells and two KO clones (sgKD5/1 and sgKD5/2) and the same cells reconstituted with the indicated FASTKD5-3xFLAG protein variants. (**A**) Indicated variants rescue the expression of COX I and cyt b and are denoted as dispensable amino acids.

WT- wild-type FASTKD5 (**B**) Indicated variants do not rescue the expression of COX I and are denoted as essential amino acids. Note, that some of these variants rescue the expression of cyt b. The three variants in purple (D249A, E317A and Y125C) are expressed at much lower level than all other variants and play a role in the stability of FASTKD5 protein. Molecular weight markers (in kDa) are indicated on the left.

Supplementary Figure S4

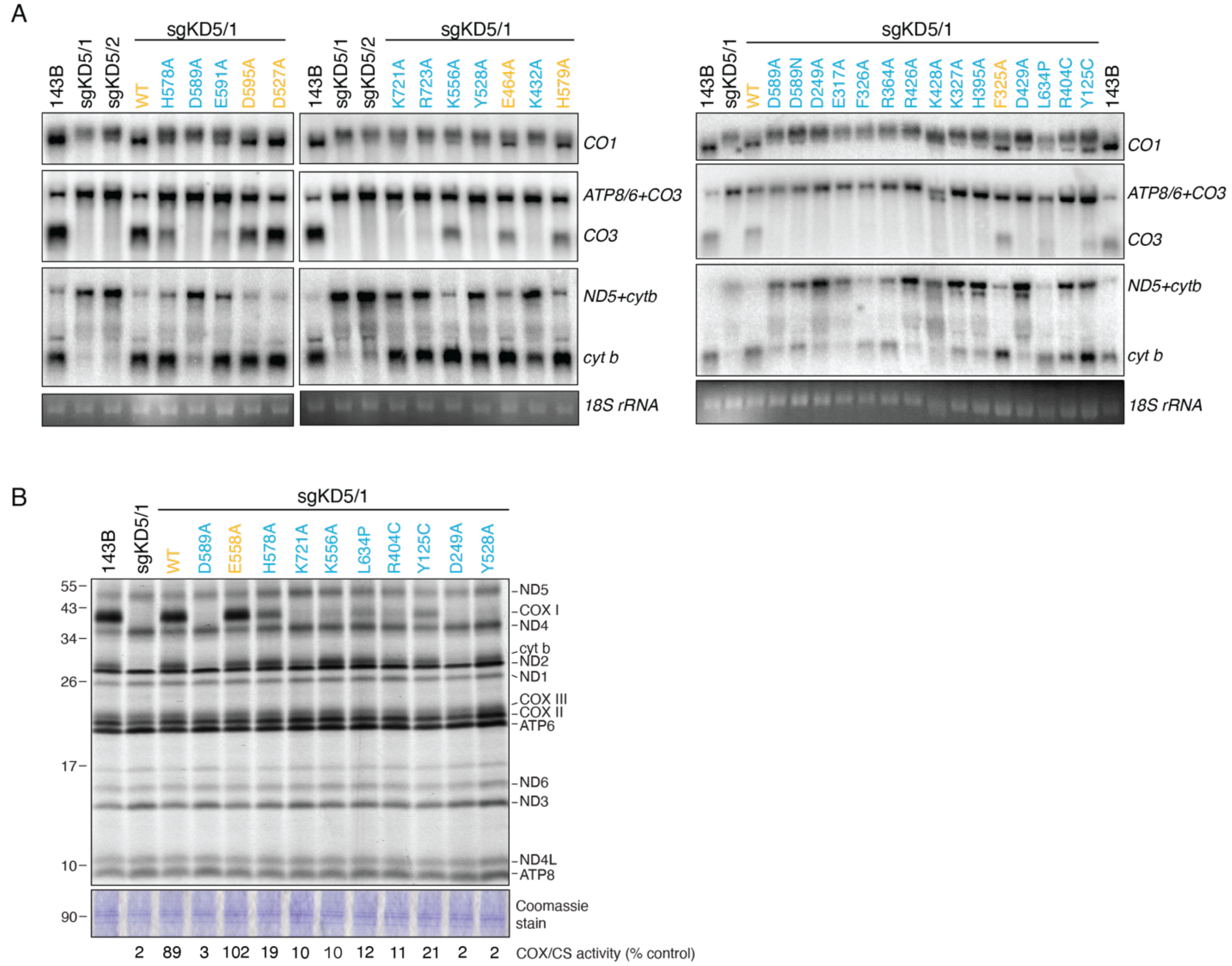

**Supplementary Figure S4** (related to **Figure 2**)

(**A**) Processing of non-canonical transcripts in 143B cells and two KO clones (sgKD5/1 and sgKD5/2) and in sgKD5/1 cells reconstituted with essential and dispensable variants for COX I expression was assessed by Northern blot analysis. UV-stain of 18S rRNA was used as a control. Image of variants K556A and Y528A plus the corresponding 143B, sgKD1 and sgKD2 are identical in Figure 2E. (**B**) Mitochondrial translation assay in selected cells confirming the lack of synthesis of COX I and cyt b polypeptides. A Coomassie total protein staining served as a loading control. COX/CS activity was measured in individual samples and is indicated as a % control (143B cells) under the gel.

Supplementary Figure S5

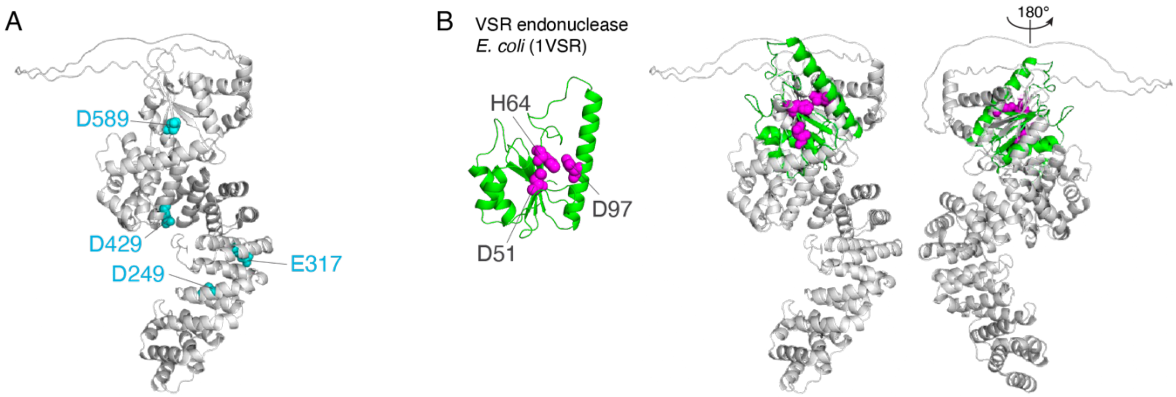

**Supplementary Figure S5** (related to **Figures 2 and 3**)

(**A**) Modelling of aspartate/glutamate residues essential for processing of all three non-canonical pre-mRNAs on Alphafold predicted FASTKD5 structure. (**B**) Modelling of VSR endonuclease from *E. coli* (1VSR, in green) on Alphafold predicted FASTKD5 structure. The active site VSR endonuclease residues are indicated in magenta.

### Supplementary Table S1

| REAGENTS | SOURCE | IDENTIFIER |
|---|---|---|
| **Antibodies** | | |
| FASTKD5 | Sigma | Cat# SAB2700438 |
| COXI | Abcam | Cat# ab14705<br>RRID: AB_2084810 |
| cyt b | Proteintech | Cat# 55090-1-AP<br>RRID:AB_2881266 |
| ND1 | a kind gift of Anne Lombes | |
| ATP6 | Proteintech | Cat# 55313-1-AP<br>RID:AB_2881305 |
| SDHA | Abcam | Cat# ab168536<br>RID:AB_2857979 |
| NDUFA9 | Abcam | Cat# ab55521<br>RID:AB_2150762 |
| ATP5A1 | Abcam | Cat# ab110273<br>RID:AB_10858175 |
| UQCRC1 | Abcam | Cat# ab110252<br>RID:AB_10863633 |
| COX IV | Abcam | Cat# ab110261<br>RRID:AB_10862101 |
| FLAG | Sigma | Cat# F1804<br>RRID:AB_262044 |
| PRDX3 | in house | |
| POLRMT | Thermo Fisher Scientific | Cat# PA5-28196<br>RRID:AB_2545672 |
| MRPP1 (TRMT10C) | Proteintech | Cat# 29087-1-AP<br>RID:AB_2881239 |
| ELAC2 | Proteintech | Cat# 10071-1-AP<br>RID:AB_2096551 |
| GRSF1 | Sigma | Cat# HPA036985<br>RID:AB_10672785 |
| DHX30 | Abcam | Cat# ab85687<br>RRID:AB_1860273 |
| TBRG4 (FASTKD4) | Sigma | Cat# HPA020582<br>RRID:AB_1857804 |
| FASTKD2 | Proteintech | Cat# 17464-1-AP<br>RID:AB_2101119 |
| MTERF3 (MTERFD1) | Sigma | Cat# HPA002966<br>RRID:AB_2147359 |
| RPUSD4 | Sigma | Cat# HPA039689<br>RRID:AB_10673537 |
| NGRN | Proteintech | Cat# 14885-1-AP<br>RID:AB_2878090 |
| RCC1L (WBSCR16) | Proteintech | Cat# 13796-1-AP<br>RID:AB_2214934 |
| GFM1 | in house | |
| TUFM/TSFM | a kind gift of Linda Spremulli | |

| | | |
|---|---|---|
| MRPS18B | Proteintech | Cat# 16139-1-AP RID:AB_2146368 |
| MRPL11 | Sigma | Cat# HPA057685 |
| LRPPRC | in house | |
| SLIRP | Abcam | Cat# ab51523 RID:AB_2066704 |
| VDAC1 | Abcam | Cat# ab14734 RRID:AB_443084 |
| DNA | Millipore | Cat# CBL186 RID:AB_11213573 |
| TRUB2 | Proteintech | Cat# 19891-1-AP RRID:AB_10640900 |
| Actin | GenScript | Cat# A00702, RRID:AB_914102 |
| Peroxidase-AffiniPure Goat Anti-Mouse IgG (H+L) antibody | Jackson ImmunoResearch Labs | Cat# 115-035-146 RRID:AB_2307392 |
| Peroxidase-AffiniPure Goat Anti-Rabbit IgG (H+L) antibody | Jackson ImmunoResearch Labs | Cat# 111-035-003 RRID:AB_2313567 |
| Goat Anti-Mouse IgG (H+L) Highly Cross-adsorbed Antibody, Alexa Fluor™ 488 Conjugated | Thermo Fisher Scientific | Cat# A-11029 RRID:AB_138404 |
| Goat anti-Rabbit IgG (H+L) Highly Cross-Adsorbed Secondary Antibody, Alexa Fluor™ 594 | Thermo Fisher Scientific | Cat# A-11037 RRID:AB_2534095 |
| Goat anti-Mouse IgG2a Cross-Adsorbed Secondary Antibody, Alexa Fluor™ 594 | Thermo Fisher Scientific | Cat# A-21135 RRID:AB_2535774 |
| Donkey Anti-Rabbit IgG (H+L) Polyclonal Antibody, Alexa Fluor™ 647 Conjugated | Thermo Fisher Scientific | Cat# A-31573 RRID:AB_2536183 |
| Goat Anti-Mouse IgG1 Antibody, Alexa Fluor™ 488 Conjugated | Thermo Fisher Scientific | Cat# A-21121 RRID:AB_2535764 |
| **Recombinant Proteins** | | |
| recombinant FASTKD5 protein | this paper | |
| **Recombinant DNA** | | |
| FASTKD5 (wild-type and all mutants) in pBABE-3xFLAG-puro | this paper | |
| pBABE-3xFLAG-Puro-gtw | in house | Aaltonen et al., LSA 2021 |
| pSpCas9(BB)-2A-Puro (PX459) V2.0 | Addgene | Cat# 62988 RRID:Addgene_62988 |
| FASTKD5 in pDONR™221 | in house | Antonicka, Cell Metab 2020 |
| pFastBac His6 TEV cloning vector with BioBrick PolyPromoter LIC Subcloning (438-B) | Addgene | Cat# 55219 RRID:Addgene_55219 |
| **Experimental Models: Cell Lines** | | |

| | | |
|---|---|---|
| 143B | ATCC | CRL-8303 |
| Phoenix | a gift from Garry P. Nolan | |
| **Oligonucleotides/oligoribonucleotides** | | |
| sgKD5-F | caccgcagtccgaagtgtgtcatac | |
| sgKD5-R | aaacgtatgacacacttcggactgc | |
| FASTKD5-E558A-a1673c | gaaggatatgaattcaaagcctgcattcttagaaactgtctttttac | |
| FASTKD5-E558A-a1673c_antisense | gtaaaaagacagtttctaagaatgcaggctttgaattcatatccttc | |
| FASTKD5-H578A-c1732g_a1733c | ggccccagtacgtcaaggcccatatgattttgcctc | |
| FASTKD5-H578A-c1732g_a1733c_antisense | gaggcaaaatcatatgggccttgacgtactggggcc | |
| FASTKD5-N604A-a1810g_a1811c | tgatgttaacctgaagccattaccatttgctagagaagccacgcc | |
| FASTKD5-N604A-a1810g_a1811c_antisense | ggcgtggcttctctagcaaatggtaatggcttcaggttaacatca | |
| FASTKD5-H579A-c1735g_a1736c | gccccagtacgtcaagcacgctatgattttgcctcatacc | |
| FASTKD5-H579A-c1735g_a1736c_antisense | ggtatgaggcaaaatcatagcgtgcttgacgtactggggc | |
| FASTKD5-D589A-a1766c | tcatacccgatcttctgccttagaggtccagcttg | |
| FASTKD5-D589A-a1766c_antisense | caagctggacctctaaggcagaagatcgggtatga | |
| FASTKD5-E606A- a1817c | cattaccatttaatagagcagccacgccggctga | |
| FASTKD5-E606A- a1817c_antisense | tcagccggcgtggctgctctattaaatggtaatg | |
| FASTKD5-E649A-a1946c | caaaactgagtcagcgcctgggcagcagc | |
| FASTKD5-E649A-a1946c_antisense | gctgctgcccaggcgctgactcagttttg | |
| FASTKD5-K696A-a2086g_a2087c | atgcagaccccaagaatggcgctggctgttcagttcac | |
| FASTKD5- K696A-a2086g_a2087c_antisense | gtgaactgaacagccagcgccattcttggggtctgcat | |
| FASTKD5-R712A-a2134g_g2135c | agtattgctatggctccgcggatctccttggactgc | |
| FASTKD5- R712A-a2134g_g2135c_antisense | gcagtccaaggagatccgcggagccatagcaatact | |
| FASTKD5-K721A-a2161g_a2162c | ccttggactgcacaatatggcgaggcggcagctg | |
| FASTKD5-K721A-a2161g_a2162c_antisense | cagctgccgcctcgccatattgtgcagtccaagg | |
| FASTKD5-R723A-c2167g_g2168c | cacaatatgaagagggcgcagctggctcggct | |
| FASTKD5- R723A-c2167g_g2168c_antisense | agccgagccagctgcgccctcttcatattgtg | |
| FASTKD5-E734A-a2201c | ggctaccgtgtggtagcgttatcctactgggaa | |
| FASTKD5-E734A-a2201c_antisense | ttcccagtaggataacgctaccacacggtagcc | |
| FASTKD5-E739A-a2216c_a2217g | gtagagttatcctactgggcgtggctcccactactgaaac | |
| FASTKD5-E739A-a2216c_a2217g_antisense | gtttcagtagtgggagccacgcccagtaggataactctac | |
| FASTKD5-E750A-a2249c_a2250g | tactgaaacgaactcgcttagcgaagttggcgtttcttcatgag | |
| FASTKD5-E750A-a2249c_a2250g_antisense | ctcatgaagaaacgccaacttcgctaagcgagttcgtttcagta | |
| FASTKD5-E757A-a2270c | agttggcgtttcttcatgcgaaagtattcacctctgc | |
| FASTKD5-E757A-a2270c_antisense | gcagaggtgaatactttcgcatgaagaaacgccaact | |
| FASTKD5-K758A-a2272g_a2273c | gttggcgtttcttcatgaggcagtattcacctctgctctc | |
| FASTKD5-K758A-a2272g_a2273c_antisense | gagagcagaggtgaatactgcctcatgaagaaacgccaac | |

| | |
|---|---|
| FASTKD5-E591A | ccgatcttctgacttagcggtccagcttgatgtta |
| FASTKD5-E591A-antisense | taacatcaagctggaccgctaagtcagaagatcgg |
| FASTKD5-D595A | cttagaggtccagcttgctgttaacctgaagccat |
| FASTKD5-D595A-antisense | atggcttcaggttaacagcaagctggacctctaag |
| FASTKD5-N604A | tgatgttaacctgaagccattaccatttgctagagaagccacgcc |
| FASTKD5-N604A-antisense | ggcgtggcttctctagcaaatggtaatggcttcaggttaacatca |
| FASTKD5-E656A | ggcagcagcccatggcgttggagaataaggc |
| FASTKD5-E656A-antisense | gccttattctccaacgccatgggctgctgcc |
| FASTKD5-E647A-a1940c | ccagggcaaaactgcgtcagagcctgggc |
| FASTKD5-E647A-a1940c-antisense | gcccaggctctgacgcagttttgccctgg |
| FASTKD5-N703A-a2107g_a2108c | gctggctgttcagttcacagccaggaaccagtattgctat |
| FASTKD5-N703A-a2107g_a2108c-antisense | atagcaatactggttcctggctgtgaactgaacagccagc |
| FASTKD5-E540A-a1619c | actcaccttcagcaagcggggtctgaattgctg |
| FASTKD5-E540A-a1619c-antisense | cagcaattcagaccccgcttgctgaaggtgagt |
| FASTKD5-K556A-a1666g_a1667c | ctgtggtatttagcagagaaggatatgaattcagcgcctgaattcttagaaa |
| FASTKD5-K556A-a1666g_a1667c-antisense | tttctaagaattcaggcgctgaattcatatccttctctgctaaataccacag |
| FASTKD5-Y528A-t1582g_a1583c | agttggcattgagtgtccagatgccagaggcaatcgtctta |
| FASTKD5-Y528A-t1582g_a1583c-antisense | taagacgattgcctctggcatctggacactcaatgccaact |
| FASTKD5-D527A-a1580c | tggcattgagtgtccagcttacagaggcaatcgtc |
| FASTKD5-D527A-a1580c-antisense | gacgattgcctctgtaagctggacactcaatgcca |
| FASTKD5-E513A-a1538c | aagtttgacctccttaaggcactatataccctcgatgg |
| FASTKD5-E513A-a1538c-antisense | ccatcgagggtatatagtgccttaaggaggtcaaactt |
| FASTKD5-E488A-a1463c | ttttggagtactttcctgtagcgttaattgatttcgctctcag |
| FASTKD5-E488A-a1463c-antisense | ctgagagcgaaatcaattaacgctacaggaaagtactccaaaa |
| FASTKD5-E464A-a1391c | tcacagaaagatgcctgcattcaaccagtacccag |
| FASTKD5-E464A-a1391c-antisense | ctgggtactggttgaatgcaggcatctttctgtga |
| FASTKD5-K432A-a1294g_a1295c | tgtcgaagtaaagatgttgccgcgattctgtggtcatttggaac |
| FASTKD5-K432A-a1294g_a1295c-antisense | gttccaaatgaccacagaatcgcggcaacatctttacttcgaca |
| FASTKD5-D249A-a746c | gctccttttggtggctgctctctggaggtacttag |
| FASTKD5-D249A-a746c-antisense | ctaagtacctccagagagcagccaccaaaaggagc |
| FASTKD5-E303A-a908c | ccaggacctaatgcaaaaattggcatcattgatccttaaatatataga |
| FASTKD5-E303A-a908c-antisense | tctatatatttaaggatcaatgatgccaatttttgcattaggtcctgg |
| FASTKD5-E317A-a950c | gatttgatcaatttggaggcggttggtaccatctgtttg |
| FASTKD5-E317A-a950c-antisense | caaacagatggtaccaaccgcctccaaattgatcaaatc |
| FASTKD5-G324A-g971c | ggttggtaccatctgtttggcgttctttaaatcaagtactaa |
| FASTKD5-G324A-g971c-antisense | ttagtacttgatttaaagaacgccaaacagatggtaccaacc |
| FASTKD5-F325A-t973g_t974c | ggttggtaccatctgtttgggggcctttaaatcaagtactaatctct |
| FASTKD5-F325A-t973g_t974c-antisense | agagattagtacttgatttaaaggcccccaaacagatggtaccaacc |
| FASTKD5-F326A-t976g_t977c | tggtaccatctgtttggggttcgctaaatcaagtactaatctctctg |
| FASTKD5-F326A-t976g_t977c-antisense | cagagagattagtacttgatttagcgaaccccaaacagatggtacca |
| FASTKD5-K327A-a979g_a980c | ggtaccatctgtttggggttctttgcatcaagtactaatctctctgaat |

| | |
|---|---|
| FASTKD5-K327A-a979g_a980c-antisense | attcagagagattagtacttgatgcaaagaaccccaaacagatggtacc |
| FASTKD5-R364A-c1090g_g1091c | ccttagtgaatattgttaaaatgttcgctttcactcacgtggatcac |
| FASTKD5-R364A-c1090g_g1091c-antisense | gtgatccacgtgagtgaaagcgaacattttaacaatattcactaagg |
| FASTKD5-H395A-c1183g_a1184c | ggagttcaaggtgtcatggccctgactctttactgctc |
| FASTKD5-H395A-c1183g_a1184c-antisense | gagcagtaaagagtcagggccatgacaccttgaactcc |
| FASTKD5-R426A-c1276g_g1277c | tcctagagtggcacactgtgcaagtaaagatgttgccaag |
| FASTKD5-R426A-c1276g_g1277c-antisense | cttggcaacatctttacttgcacagtgtgccactctagga |
| FASTKD5-K428A-a1282g_a1283c | tcctagagtggcacactgtcgaagtgcagatgttgccaag |
| FASTKD5-K428A-a1282g_a1283c-antisense | cttggcaacatctgcacttcgacagtgtgccactctagga |
| FASTKD5-D429A-a1286c | cacactgtcgaagtaaagctgttgccaagattctgtg |
| FASTKD5-D429A-a1286c-antisense | cacagaatcttggcaacagctttacttcgacagtgtg3 |
| FASTKD5-K361A-a1081g_a1082c-new | gtcgctccttagtgaatattgttgcaatgttccgtttcactcacgtg |
| FASTKD5-K361A-a1081g_a1082c-antisense-new | cacgtgagtgaaacggaacattgcaacaatattcactaaggagcgac |
| FASTKD5-L634P-t1901c | agatgatttgatgaataagttaccaaaagggaaagcaagaggacatt |
| FASTKD5-L634P-t1901c-antisense | aatgtcctcttgctttcccttttggtaacttattcatcaaatcatct |
| FASTKD5-R404C-c1210t | ctttactgctcggccttatgcttcctgaatgaag |
| FASTKD5-R404C-c1210t-antisense | cttcattcaggaagcataaggccgagcagtaaag |
| FASTKD5-Y125C-a374g | tttcctacagctaagaccagaatgccgtgttcacag |
| FASTKD5-Y125C-a374g-antisense | ctgtgaacacggcattctggtcttagctgtaggaaa |
| FASTKD5-H220Y-c658t | gaaagcttttgtcattttaggaatcccttactcccattcaatgc |
| FASTKD5-H220Y-c658t-antisense | gcattgaatgggagtaagggattcctaaaatgacaaaagctttc |
| FASTKD5-H234W-c700t_a701g_t702g | tgtgtatgagaccaagtgttgctggcaggtatgggagatgaatatgg |
| FASTKD5-H234W-c700t_a701g_t702g-antisense | ccatattcatctcccatacctgccagcaacacttggtctcatacaca |
| FASTKD5-I264Y-a790t_t791a | gccgcaaagtacctaggtttttaaactatttttctagttatcttaatttgcact |
| FASTKD5-I264Y-a790t_t791a-antisense | agtgcaaattaagataactagaaaaatagtttaaaaacctaggtactttgcggc |
| FASTKD5-R500W-a1498t | cagtccagggtttgtctggttagctcaggagag |
| FASTKD5-R500W-a1498t-antisense | ctctcctgagctaaccagacaaaccctggactg |
| FASTKD5-S221I-t661a_c662t | ttgtcattttaggaatccctcacatccattcaatgctagatgtgtatg |
| FASTKD5-S221I-t661a_c662t-antisense | catacacatctagcattgaatggatgtgagggattcctaaaatgacaa |
| FASTKD5-S223E-t667g_c668a | ttaggaatccctcactcccatgaaatgctagatgtgtatgagac |
| FASTKD5-S223E-t667g_c668a-antisense | gtctcatacacatctagcatttcatgggagtgagggattcctaa |
| FASTKD5-S267Q-a799c_g800a_t801a | cgcaaagtacctaggtttttaaacatttttctcaatatcttaatttgcactggaaggatcta |
| FASTKD5-S267Q-a799c_g800a_t801a-antisense | tagatccttccagtgcaaattaagatattgagaaaaatgtttaaaaacctaggtactttgcg |
| FASTKD5-W237A-t709g_g710c | ccaagtgttgccatcaggtagcggagatgaatatggatca |
| FASTKD5-W237A-t709g_g710c-antisense | tgatccatattcatctccgctacctgatggcaacacttgg |
| FASTKD5-W237Y-g710a_g711t | ccaagtgttgccatcaggtatatgagatgaatatggatcagct |

| | |
|---|---|
| FASTKD5-W237Y- g710a_g711t-antisense | agctgatccatattcatctcatatacctgatggcaacacttgg |
| FASTKD5-noRAP | cagaccccaagaatgaaggacccagctttcttgtac |
| FASTKD5-noRAP-antisense | gtacaagaaagctgggtccttcattcttggggtctg |
| FASTKD5-noFAST_1 | cgaagtaaagatgttgccaggttagctcaggagaga |
| FASTKD5-noFAST_1-antisense | tctctcctgagctaacctggcaacatctttacttcg |
| FASTKD5-noFAST_2 | gaactaagtttgacctcctttttaatagagaagccacgcc |
| FASTKD5-noFAST_2-antisense | ggcgtggcttctctattaaaaaggaggtcaaacttagttc |
| FASTKD5-delta2-27 | agcaggctccaccatggtgtcatactggaatg |
| FASTKD5-delta2-27-antisense | cattccagtatgacaccatggtggagcctgct |
| FASTKD5-delta36-420 | ctggaatgtgagcagcagagtggcacactgtc |
| FASTKD5-delta36-420-antisense | gacagtgtgccactctgctgctcacattccag |
| *5'end-CO1-3'Cy3* | rArUrUrUrUrArCrCrUrCrArCrCrCrCrCrArCrUrGrArUrGrUrUrCrGrCrCrGrArCrCrGrUrUrGrArCrUrA |
| *ATP6-CO3-3'Cy3* | rCrUrGrCrArCrGrArCrArArCrArCrArUrArArUrGrArCrCrCrArCrCrArArUrCrArCrArUrGrCrCrUrArU |
| *ND5-cytb-3'Cy3* | rCrArArCrUrArCrArArGrArArCrArCrCrArArUrGrArCrCrCrCrArArUrArCrGrCrArArArArCrUrArArC |
| *CO1_309-349-3'Cy3* | rArCrUrCrUrUrArCrCrUrCrCrCrUrCrUrCrUrCrCrUrArCrUrCrCrUrGrCrUrCrGrCrArUrCrUrGrCrUrA |
